## Supplementary Materials for "Emotionotopy in the Human Right Temporo-Parietal Cortex"

Lettieri et al.

- **Supplementary Methods pages 2-13**


*Generalization of emotion dimensions to a wider sample of emotion categories* pag.2
*Does Forrest Gump reflect real life emotion dynamics?* pag.3
*fMRI data pre-processing* pag.5
*Searchlight analysis* pag.6
*Effect size and noise-ceiling estimation* pag.7
*Rotation of the emotion dimension model* pag.8
*Comparison between emotion gradients and meta-analytic definition of right TPJ* pag.11
*Do emotion dimension gradients depend on low-level acoustic and visual features?* pag.12


- **Supplementary Figures pages 14-30**


*SF1. Brain regions encoding emotion ratings* pag.14
*SF2. Characterization of emotion gradients* pag.15
*SF3. Single-subject emotion gradients in right TPJ* pag.16
*SF4. Testing rotated versions of emotion dimensions in right TPJ* pag.17
*SF5. Brain areas representing the combination of emotion dimension gradients as revealed by searchlight analysis* pag.18
*SF6. Brain areas representing individual emotion dimension gradients as revealed by searchlight analysis* pag.19
*SF7. Emotion dimension gradients using unsmoothed fMRI data* pag.20
*SF8. Subjective emotion rating model versus third person emotion attribution models* pag.21
*SF9. Preferred responses of distinct populations of voxels using NNMF* pag.22
*SF10. Fitting of low-level features* pag.23
*SF11. PCs loadings obtained from the emotion attribution model* pag.24
*SF12. Real life versus Forrest Gump emotion transitions* pag.25
*SF13. fMRI data single-subject preprocessing* pag.26
*SF14. fMRI data group-level preprocessing* pag.27
*SF15. Comparison between emotion gradients and meta-analytic definition of right TPJ* pag.28
*SF16. Regions associated to the direction of portrayed emotions* pag.29
*SF17. Emotion dimension gradients do not depend on low-level acoustic and visual features* pag.30


- **Supplementary Tables pages 31-36**


*ST1. Brain regions encoding emotion ratings* pag.31
*ST2. Emotion gradients in TPJ* pag.32
*ST3. Emotion gradients in TPJ relative to emotion dimensions or basic emotions* pag.33
*ST4. Single-subject emotion dimension gradients in right TPJ* pag.34
*ST5. Topographic organization of portrayed emotions in right TPJ* pag.35
*ST6. Topographies in right TPJ considering spatial smoothing and cortical folding* pag.36


- **Supplementary References pages 37-38**

### Supplementary Methods

#### Generalization of emotion dimensions to a wider sample of emotion categories

To test whether our emotion dimensions depended on behavioral ratings of six basic emotions, we verified their existence starting from a comprehensive set of emotion categories, which included secondary affective states as well. We employed Labs and colleagues[1] ratings describing portrayed emotions of Forrest Gump characters considering embedded affective states (i.e., other-directed emotions). Participants tagged 205 randomly presented movie segments choosing among a large set of emotion categories[2] (N=22) and were allowed to watch each scene more than once. Other than happiness, fear, sadness and anger, these 22 emotion categories included secondary and social states as admiration, contempt, gratitude, hate, love, and pride among others (for a complete description please refer to[1]). Thus, we applied Principal Component (PC) analysis to Labs data after lagging and temporally smoothing the 22 emotion timeseries, as we did for the subjective emotion ratings. The first six dimensions (~85% of explained variance) were selected to match the dimensionality of our emotion rating model and were transformed by rotating PC scores using the procrustes criterion. Results of this procedure are presented in Supplementary Figure 11, in which factor loadings of polarity, complexity and intensity dimensions (panel A) can be compared with the unrotated (panel B) and rotated (panel C) version of Labs and colleagues[1] PCs. The first three rotated components represented respectively the 20.6%, 19.8% and 16.6% of the explained variance, and were positively associated with our three emotion dimensions (panel E). Correlation for rotated $PC_1$ versus polarity was Spearman's $\rho = 0.589$, for rotated $PC_2$ versus complexity was Spearman's $\rho = 0.533$ and for rotated $PC_3$ versus intensity was Spearman's $\rho = 0.488$.

It is important to note that, other than basic emotions (i.e., happiness, fear, sadness and anger), only four secondary/social affective states - i.e., love, contempt, admiration and gloating - substantially contributed to the first six components derived from Labs and colleagues data, even considering the unrotated version (panel B). Indeed, the majority of emotional episodes involved the five categories of anger, fear, happiness, love and sadness, whereas other secondary/social categories available to subjects (e.g., resentment, gratification, satisfaction) were used infrequently or employed only by a subset of observers[1].

In summary, the same polarity, complexity and intensity dimensions emerge even when a broader set of emotion categories are used.

### Does Forrest Gump reflect real life emotion dynamics?

Forrest Gump is an emotionally evocative movie that elicits a variety of affective states in a relatively short amount of time. Although movies have been successfully used to study emotions in the laboratory setting[3-5], we cannot exclude that the dynamics of portrayed emotions mimic those experienced in the real life. To explore this possibility, we took advantage of Thornton & Tamir experience-sampling dataset[6] (i.e., Study 3, see also[7]) comprising ~65,000 ratings obtained from ~10,000 participants, who were asked to report their own emotional state throughout the day, choosing among 18 categories (i.e., alertness, amusement, awe, gratitude, hope, joy, love, pride, satisfaction, anger, anxiety, contempt, disgust, embarrassment, fear, guilt, offense and sadness). In this study, the authors used the collected reports to build an experience-based description of emotion transitions (i.e., real life emotion transitions). Specifically, by considering each reported emotion and the one following in time, they tested whether the co-occurrence of emotions is predicted by a mental representation of emotion transitions (for further details please refer to[6]). We particularly selected the model based on study 3[6], as nine out of 18 emotion categories included in this dataset (i.e., anger, sadness, fear, contempt, satisfaction, gratitude, hope, love and pride) were also adopted by Labs and colleagues[1] to label portrayed emotions in Forrest Gump. Starting from these data, we thoroughly followed the methods reported in[6] and converted ratings into discrete outcomes (i.e., emotion present or not) for each timepoint. We then built a transition count matrix by measuring the number of transitions between all possible emotion pairings in adjacent timepoints (i.e., between t and t+1). This matrix was further normalized by frequency-based expectations obtaining the odds of each transition. The log-transformed version of this matrix (i.e., movie emotion transitions) was then compared to real-life data using Spearman's ρ. To assess the statistical significance of this association, we generated surrogate timeseries for the nine emotion categories through the IAAFT procedure (N = 1,000; see Methods in the main manuscript for details). For each of the 1,000 null models, a transition count matrix was then obtained, normalized and log-transformed. The obtained matrices were correlated with real-life data, generating a null distribution against which the actual association between movie and real life emotion transitions was tested.

Results showed that emotion transitions obtained from movie and real-life data were significantly associated (Spearman's $\rho = 0.646$; $p = 0.001$; Supplementary Figure 12). In addition, as this analysis explores the similarity between movie and real life data in a short time window (2s), we also evaluated whether this relationship exists at different time scales. Therefore, we built a number of movie-based models, each measuring the likelihood of emotion transitions between timepoint t and timepoint t+n in the future, with a maximum delay of 120 seconds (60 timepoints). These models were then correlated with real-life data and statistical significance was assessed using the procedure described above. Results are reported in panel D of Supplementary Figure 12 and show that the real life model predicts emotion transitions in the movie up to 58 seconds.

Of note, happiness is one of the emotion categories most present in Forrest Gump tagging data, yet it has not been used in reports collected for study 3[6]. Hence, we decided to include this emotion in the movie model using joy, awe or amusement as its counterpart in the real life model. This allowed us to estimate the robustness of the association between movie and real-life data considering different facets of the basic emotion happiness.

Interestingly, using joy, awe or amusement as proxies of happiness, the association between movie and real life emotion transitions is significant (joy: Spearman's $\rho = 0.702$; $p = 0.001$; awe: Spearman's $\rho = 0.702$; $p = 0.001$; amusement: Spearman's $\rho = 0.686$; $p = 0.001$). In all these three cases, emotion transitions observed in real-life data predict those occurring in the movie up to 64 seconds in the future.

Altogether, these analyses show that within a ~60 seconds time window our stimulus reflects emotion transitions similar to those experienced in real life and predicted by a mental model of emotion co-occurrence. These findings substantiate the ecological validity of our stimulus.

### fMRI data pre-processing

We employed ANTs[8] and AFNI[9] v.17.2.00 to preprocess MRI data (Supplementary Figure 13). First, structural images were brain extracted (antsBrainExtraction.sh) and non-linearly transformed to match the MNI152 template (3dQwarp). The estimated deformation field was subsequently used to bring single-subject activation maps from the original to the standard space. Functional data were corrected for intensity spikes (3dDespike) and adjusted for slice timing acquisition (3dTshift). We also compensated head movements by registering each volume to the most stable timepoint (3dvolreg). In this regard, a rigid body transformation was adopted and the six estimated motion parameters were included as confounds in further analyses. The transformation matrices were also used to compute an aggregated measure - framewise displacement[10] - that highlighted timepoints affected by excessive motion. Functional data were linearly (align_epi_anat.py) and non-linearly registered to the T1w images, also correcting for phase distortion, and warped to match the MNI152 template using the already computed deformation field (3dNwarpApply). Furthermore, timeseries were smoothed until they reached a full width at half maximum of 6mm (Gaussian kernel). In this regard, we did not simply apply a 6mm smoothing filter to the original data, rather we adopted the AFNI's 3dBlurToFWHM tool, which estimates and iteratively increases the smoothness of data until a specific FWHM level is reached. Lastly, we ruled out the effects of signal drifts, head motion and heartbeat (3dDeconvolve) to obtain timeseries of brain activity cleaned from these nuisance regressors.

Following the same procedure adopted for the behavioral processing, single-subject preprocessed fMRI data were averaged to obtain group-level hemodynamic activity and for each voxel the same windowing procedure was employed to temporally smooth data (moving average: 10s window;

Supplementary Figure 14). From the obtained aggregated and smoothed timeseries, the timecourse of low-level acoustic (i.e., volume energy - RMS of the signal) and visual (i.e., Gabor contrast energy for 0.5 and 8 cyc/deg spatial frequencies for each frame) features of the movie were regressed out to mitigate the possible collinearities between emotion ratings and psychophysical properties of the stimulus (e.g., fearful events might be associated to sudden volume increases). Specifically, the RMS value was estimated on 2s non-overlapping windows[11] matching the TR of the fMRI scan. For the low-level visual features instead, we modeled the canonical response of area V1[12]. Each movie frame was filtered with a set of oriented Gabor filters encompassing the lowest and highest limits of V1 spatial frequency selectivity (0.5 and 8 cyc/deg), as found by cell recordings in non-human[13] and by fMRI in humans primates[14,15]. Filters response was averaged across four orientations (i.e., 0, 45, 90, 135 deg) and all pixels, to obtain a global descriptor for each frequency in each frame. Visual features were then temporally averaged across frames, delayed and smoothed in time to match the temporal resolution of fMRI data.

Overall, low-level features modelling generated three regressors of no interest (i.e., low and high spatial frequencies of movie frames and RMS of the audio track) that were regressed out from brain activity using a multiple regression analysis. Results for the fitting of low-level features are depicted in Supplementary Figure 10. The obtained regression residuals, consisting of 3,595 timepoints, were considered as the dependent variable in the encoding analysis having emotional ratings as predictors.

### Searchlight analysis

We performed a data-driven searchlight analysis to test whether right TPJ was the only region significantly encoding all the three emotion dimension gradients. Thus, for each voxel significantly associated to emotion ratings (i.e., shaded and outlined regions in Supplementary Figure 5) we built a spherical region of interest (i.e., searchlight; 15mm radius) and derived the Euclidean distance of voxel coordinates and of $\beta$ coefficients related to the fitting of the three emotion dimensions.

Functional and anatomical dissimilarity matrices were then compared using Spearman's ρ coefficient and the computation of p-value was based on surrogate data (i.e., 1,000 IAAFT-based null models). Results were corrected using the False Discovery Rate procedure and minimum cluster size > 10. The combination of the three emotion dimension gradients was represented within a patch of cortex centered in right pSTS/TPJ only (Supplementary Figure 5; red-colored regions; $q < 0.05$ FDR corrected; CoG: x = 58, y = -53, z = 21). This evidence corroborated the original findings based on the hypothesis-driven approach (i.e., NeuroSynth "TPJ").

Furthermore, we searched for individual emotion dimension topographies in regions encoding the emotion rating model. To do this, we ran three separate searchlight analyses measuring the topographic arrangement of polarity, complexity and intensity. The resulting maps were then combined into a comprehensive description of the distribution of gradients across the brain. Briefly, we employed a specific coding in the RGB color space[16]. The red channel was assigned to polarity, the green to complexity and the blue to intensity. Color brightness relates to the log transformed p-value of the fitting of each component. This procedure highlighted regions predominantly involved either in polarity, complexity or intensity, as well as in any combination of the three (Supplementary Figure 6). Results showed that right pSTS/TPJ region is the only area of overlap of the three emotion dimension gradients even considering polarity, complexity and intensity separately.

### Effect size and noise-ceiling estimation

To evaluate the effect size of the association between emotion ratings and right TPJ activity, we correlated the predicted fMRI signal obtained from the encoding procedure, with the actual BOLD activity within the same peak voxel (i.e., $R^2 = 0.07$). The association between the two timeseries was Spearman's $\rho = 0.23$ and Kendall's $\tau = 0.15$.

To allow a direct and unbiased comparison between $R^2$ values obtained from the fitting of different emotion models in right TPJ, we also performed a cross-validation using a half-run split method

(Supplementary Figure 8). Specifically, we randomly selected one of the two halves as the training data for the estimation of β coefficients. We then measured the goodness of fit of our model by multiplying the predictors of the remaining half with estimated β coefficients, thus reconstructing the predicted fMRI signal. The latter was then correlated with the actual fMRI activity, obtaining the final cross-validated $R^2$ coefficient. To avoid possible confounds introduced by selecting the first or the second part of each run as training/test dataset, we repeated the same procedure 200 times (i.e., bootstrapping), each one randomly assigning the first or second half to the training/test set. The use of this procedure resulted in an effect size of $R^2 = 0.04$ for the right TPJ peak.

Moreover, we conducted a noise-ceiling analysis for right TPJ data, similarly to what has been done by Ejaz and colleagues[17]. For each right TPJ voxel, we calculated the average association (i.e., $R^2$ value) between single-subject timeseries and group-level activity. This procedure considers group-level fMRI data as the ground-truth model. However, the average signal is biased as it includes single-subject information from all the enrolled participants, ultimately producing an overestimate of the actual noise-ceiling level (i.e., the upper bound). Therefore, to obtain an estimate of the lower bound of noise-ceiling, we iteratively measured the association between each individual timeseries and the group-level average signal obtained from all the other participants (i.e., leave-one-subject-out procedure). We found that lower and upper noise ceiling bounds of the right TPJ peak voxel were $R^2 = 0.13$ (Spearman's $\rho = 0.33$ and Kendall's $\tau = 0.22$) and $R^2 = 0.23$ (Spearman's $\rho = 0.45$ and Kendall's $\tau = 0.31$), respectively.

### Rotation of the emotion dimension model

We developed a novel approach to test the correspondence between anatomo-functional gradients and PC rotations, to reveal which stimulus features are actually encoded onto the cortical mantle[18].

First, we restricted our analysis to the three emotion dimensions consistent across subjects (i.e., polarity, complexity and intensity), which showed a gradient-like organization in right TPJ as well. Second, we performed only orthogonal rotations because of two reasons: (1) any orthogonal rotation of the original components will explain the same total variance; (2) the computation of gradient direction requires the accurate estimate of β coefficients obtained from a multiple linear regression analysis. This approach is however not robust if predictors are collinear, which may be the case when oblique rotations are applied. Therefore, we first estimated all the possible elemental rotations along the axes defined by the three emotion dimensions (i.e., x: polarity, y: complexity and z: intensity). We explored rotations between ±45° with 1° step, as this range ensured univocal solutions that would not produce the shifting of PC labels. As a matter of fact, considering a convenient bi-dimensional example, we can assert that 60° orthogonal rotations for $PC_1$ and $PC_2$ would produce solutions in which $PC_1$ approximates the unrotated version of $PC_2$ and $PC_2$ resembles the 180°-rotated (i.e., flipped) version of $PC_1$. Such a solution, though, would be identical to a 30° rotation, except for the PC sign. In line with this, rotations of ±90° would simply shift PC labels (e.g., rotated complexity would become now unrotated intensity), whereas ±180° rotations would result in sign flipping. The latter case leads to brain activity estimates (i.e., β values) being the topographically mirrored version of those obtained using the unrotated dimensions and, thus, to ρ values of the same magnitude for the association between anatomical and functional distance. As all the possible rotations between ±45° produce ~750k solutions - which is already computationally intense -, we uniformly sampled 70k rotations from the original space. Further, the intuitive mapping of gradient magnitude (i.e., Spearman's ρ between anatomical and functional distance) in the manifold defined by the rotated solutions is non trivial and the method we propose is illustrated in Supplementary Figure 4A.

In brief, we represented gradient intensity of the unrotated emotion dimensions as the central point of a 3D manifold described by all the ±45° explored rotations. We also mapped gradient intensity of all the rotated solutions as points in this space, color-coding the magnitude of the association between anatomical and functional distance. Rotations are expressed according to three cardinal trajectories originating from the central point (i.e., the unrotated emotion dimensions), each one determining the orthogonal rotation of two components while maintaining fixed the other one.

Therefore, points lying on the red trajectory depict solutions in which the original unrotated version of polarity is present and complexity and intensity are actually rotated. The same applies also to the green and blue trajectories in which complexity and intensity respectively maintain their original unrotated form. All the other mapped solutions describe orthogonal rotations concurrently applied to the three emotion dimensions. The larger the geodesic distance in the solution space between axes origin and a specific point, the larger is the applied rotation to the original emotion dimensions. Lastly, the position of each solution with respect to the central point also defines the direction of the rotation (i.e., positive or negative).

Results show that the original unrotated version of the polarity, complexity and intensity dimensions is the optimal solution to explain the gradient-like organization of right TPJ. Indeed, within the space defined by PC rotations, no solutions retained ρ coefficients (i.e., gradient magnitude) larger than those associated with the unrotated components for all the three emotion dimensions.

In addition, rotations in which the gradient magnitude is similar across the three emotion dimensions are arranged close to the unrotated solution (i.e., white areas in Supplementary Figure 4B), whereas moving away from axes origin at least one of the three dimensions is not represented as a gradient in right TPJ (i.e., yellow and cyan areas in Supplementary Figure 4B). Of note, considering all the explored solutions, very few rotations produce gradients encoding combined polarity and intensity, but not complexity (i.e., lack of magenta areas in Supplementary Figure 4B).

As the original unrotated solution was the best among ~70k explored rotations, we assessed the probability of occurrence of such behavior using a Monte Carlo simulation. Therefore, we created 1,000 PC models by selecting 100 consecutive timepoints from the emotion dimension timeseries to predict randomly sampled right TPJ activity (N = 100 consecutive timepoints). For each iteration, we then mapped the results of the multiple linear regression analysis (i.e., β coefficients) on a 3-D grid of 25 voxels and computed the correspondence between the anatomical and functional distance obtained using the unrotated and rotated (±45° with 5° step; ~7k explored solutions) predictors.

Lastly, we counted the number of iterations in which the gradient magnitude of the rotated predictors was higher with respect to the original unrotated solution.

Results of the Monte Carlo simulation confirm the peculiarity of real data. Indeed, while the unrotated version of emotion dimensions represents the optimal solution in explaining right TPJ topography, rotated components produce stronger gradients in the vast majority of simulated cases (96.2%; $p < 0.05$). Of note, we tested the reliability of the results obtained from the Monte Carlo simulation by also varying the length of the timeseries (50, 100 and 200 timepoints), the number of voxels (N = 25, 100) and by generating synthetic PC models and fMRI signal using Gaussian noise. Results for all these procedures were consistent with the original simulation (data not shown).

The code for computing, exploring and rendering PC rotations is available in the online repository, as reported in the manuscript.

### Comparison between emotion gradients and meta-analytic definition of right TPJ

The existence of emotion dimension gradients generalizes across several definition of the ROI size, yet the optimal solution is represented by a 15mm radius sphere (11,556 $mm^3$ volume). In fact, although emotion dimension gradients are significantly represented also considering a 27mm ROI (i.e., Supplementary Table 2), the effect size decreases for radii larger than 15mm. To clarify the extent of our emotion dimension gradients, we performed a quantitative comparison of the size of our ROI with the definition of right TPJ based on the neuroimaging literature.

To do so, we considered the right TPJ region obtained from the Neurosynth database (http://old.neurosynth.org/analyses/terms/tpj/). This meta-analytic definition relies on brain activations elicited by classic Theory of Mind and affective processing tasks, such as false-belief[19,20], emotion perception[21] or reappraisal tasks[22], providing a reliable estimate of the right TPJ size. Considering the Neurosynth TPJ reverse inference map - i.e., p(F|A) -, the volume of the largest cluster was 8,127 $mm^3$ (coordinates: x = +58, y = -50, z = +16), whereas the volume of the spherical ROI that better represents emotion topography in our study (i.e., 15mm radius) was

11,556 $mm^3$. Yet, considering the TPJ forward inference map - i.e., p(A|F) -, the volume of the largest cluster was 16,929 $mm^3$ (coordinates: x = +58, y = -50, z = +16). Altogether, these results indicate that the optimal description of emotion dimension gradients is represented in a patch of cortex that approximates the definition of right TPJ based on brain activation studies (i.e., ~42% larger in volume as compared to the reverse inference map, but also ~32% smaller than the forward inference definition). Supplementary Figure 15 depicts a comparison between the Neurosynth map and our spherical ROI.

### Do right TPJ emotion dimension gradients depend on low-level acoustic and visual features?

To further ensure that the right TPJ emotion dimension gradients do not depend on low-level sensory information confounds, we built more complex descriptions of visual and acoustic features of Forrest Gump, based on well-established models.

Specifically, we selected spectral power density as a model of low-level acoustic information[23], and GIST descriptors for visual features[24-26]. We derived the power spectrum for each 2 s segment of the audio track and calculated the power in dB units. The procedure we used is identical to the one described in de Heer and colleagues[23]: Welch method, Gaussian window with SD of 5 ms, length 30 ms, 1 ms spacing between windows. The resulting model comprised 449 columns describing the power spectrum of the acoustic signal ranging from 0 Hz to 15 kHz in steps of 33.5 Hz.

For the visual model, we segmented each movie frame into a 4x4 grid and sampled the responses to Gabor filters having four different sizes and four possible orientations. This procedure generated a vector of 256 elements, which described each video frame in terms of spatial frequencies, Gabor filter orientations and positions in the visual field. All the GIST descriptors were averaged within a 2 s time window. Timeseries of 449 acoustic and 256 visual features were lagged by 2s and temporally smoothed using a 10s window, similarly to the emotion ratings model. As all our procedures rely on multiple linear regression, which advocate for the use of orthogonal predictors, we performed a PC analysis on the acoustic and visual models separately and selected the first 21 PCs, which explained more than 90% of the total variance. We then regressed out low-level visual and acoustic stimulus features from brain activity and tested the existence of right TPJ emotion dimension gradients. Importantly, right TPJ emotion dimension gradients were not affected by regressing out low-level properties from BOLD signal: polarity ($\rho = 0.258$, p-value = 0.031, 95% CI: 0.252 to 0.264), complexity ($\rho = 0.261$, p-value = 0.013, 95% CI: 0.254 to 0.267) and intensity ($\rho = 0.270$, p-value = 0.016, 95% CI: 0.264 to 0.277). Overall, this evidence indicates that the topographic organization of affective states in right TPJ is not explained by low-level sensory information confounds. Procedures and relative results are summarized in Supplementary Figure 17.

### Supplementary Figures

#### Supplementary Figure 1. Brain regions encoding emotion ratings

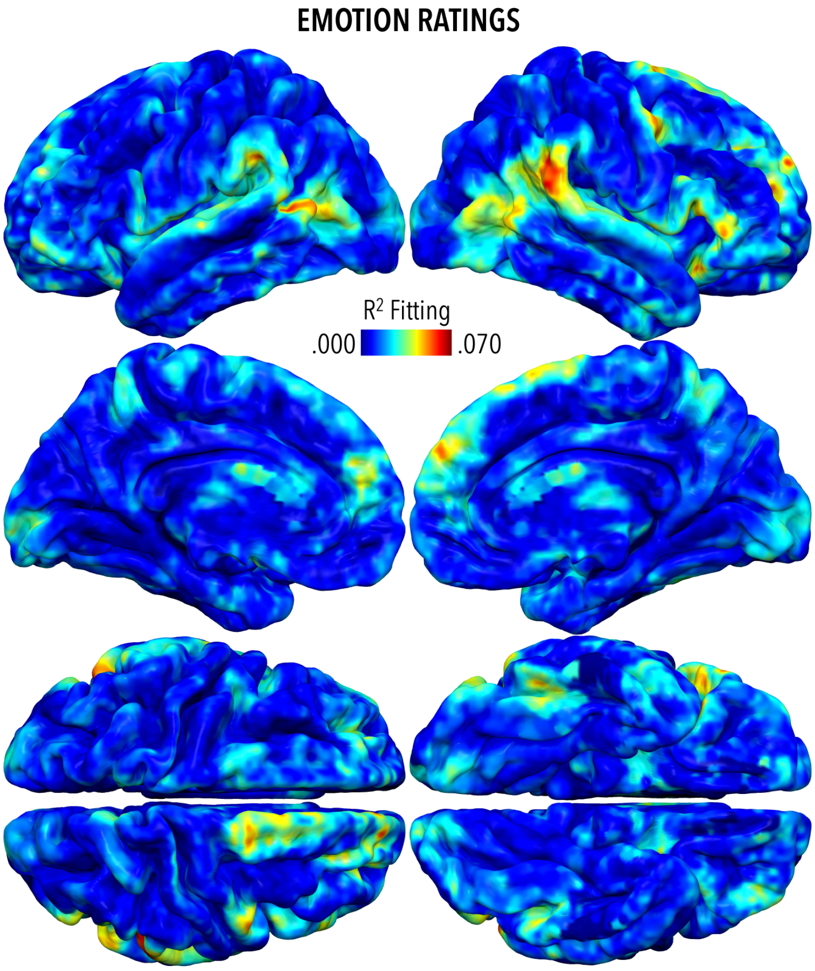


Map of the $R^2$ fitting of emotion ratings. Datasets relative to these results are available in the public repository.

### Supplementary Figure 2. Characterization of emotion gradients

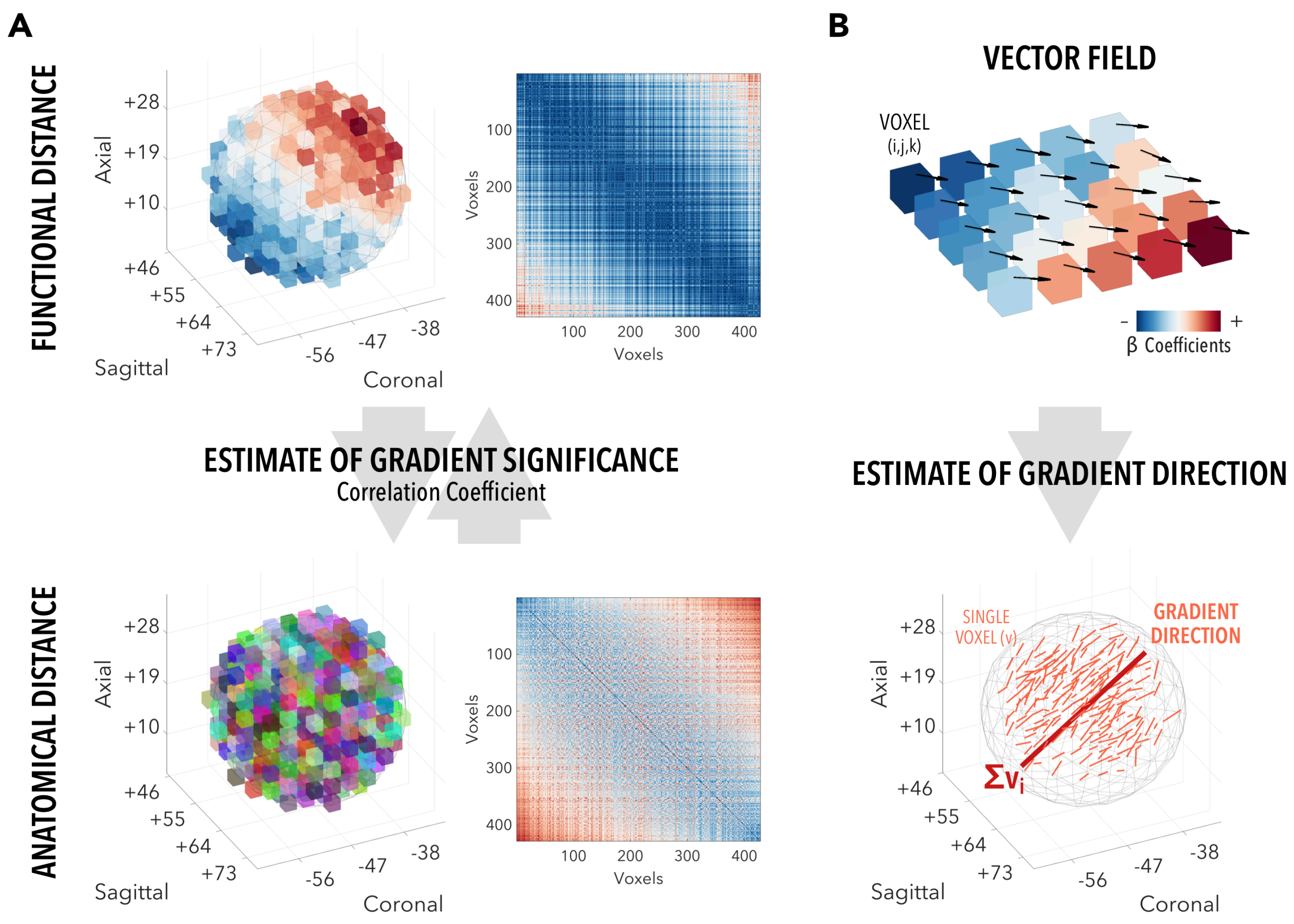


The presence of a gradient-like organization is verified by testing the similarity between functional and anatomical information. Starting from a specific patch of cortex, two dissimilarity matrices are computed (**A**): one using the Euclidean distance of voxel coordinates (i.e., anatomical distance), and the other one using the Euclidean distance of β coefficients related to the fitting of a specific model (i.e., functional distance). Spearman's ρ is used to measure the strength and assess the significance of the relationship (panel **A**). To derive the main direction of a (linear) gradient, the vector field determined by β coefficients is then estimated and summed across voxels (panel **B**).

**Supplementary Figure 3. Single-subject emotion gradients in right TPJ**

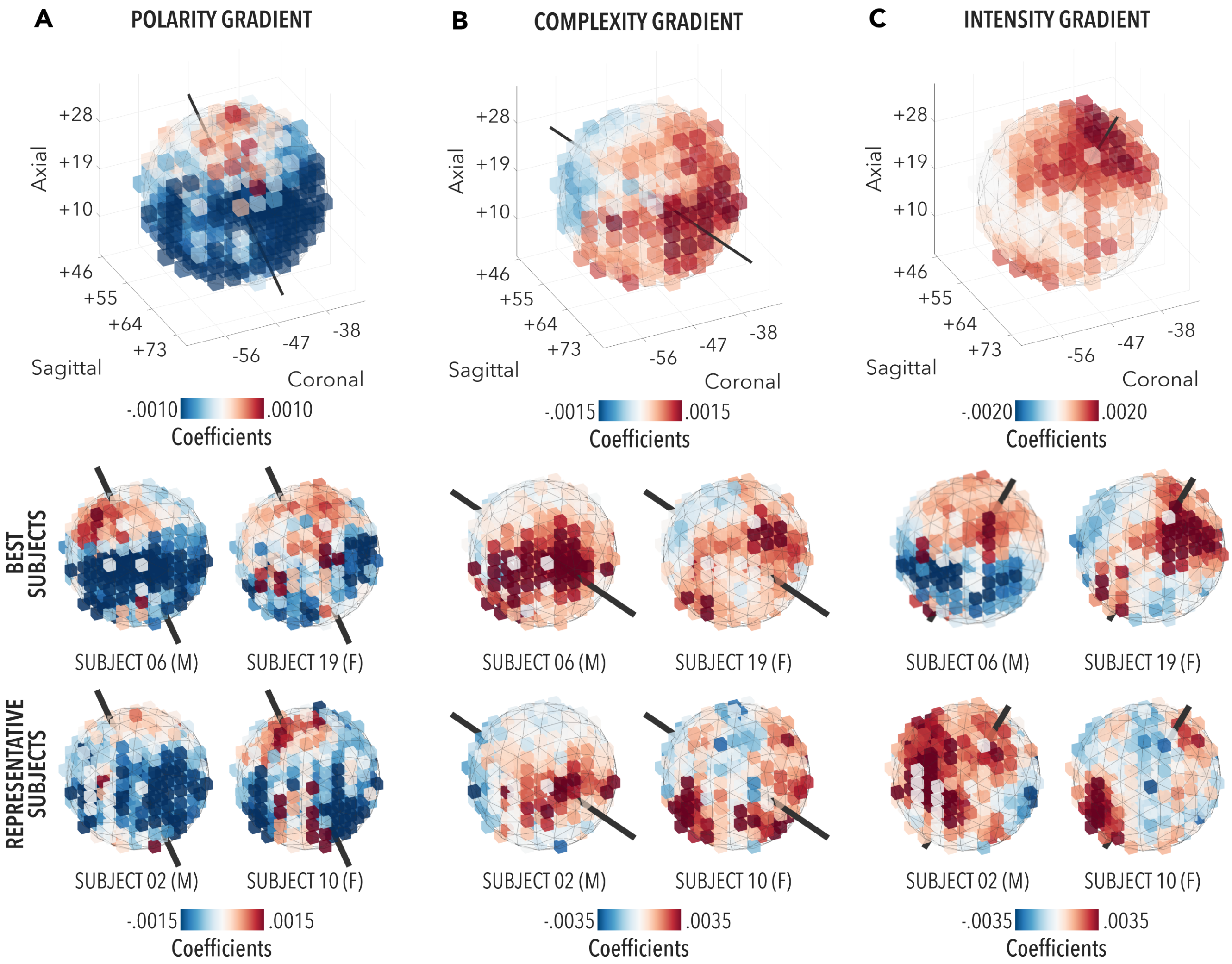


Upmost part of the figure depicts group-level results of gradient mapping (as in Figure 4). Below, results obtained from two of the best subjects (first row; one male and one female) and for two representative subjects (second row; one male and one female). Subjects coding is the one adopted in the studyforrest project. Column **A, B** and **C** report β coefficients of the polarity, complexity and intensity gradients, respectively.

### Supplementary Figure 4. Fitting rotated emotion dimensions in right TPJ

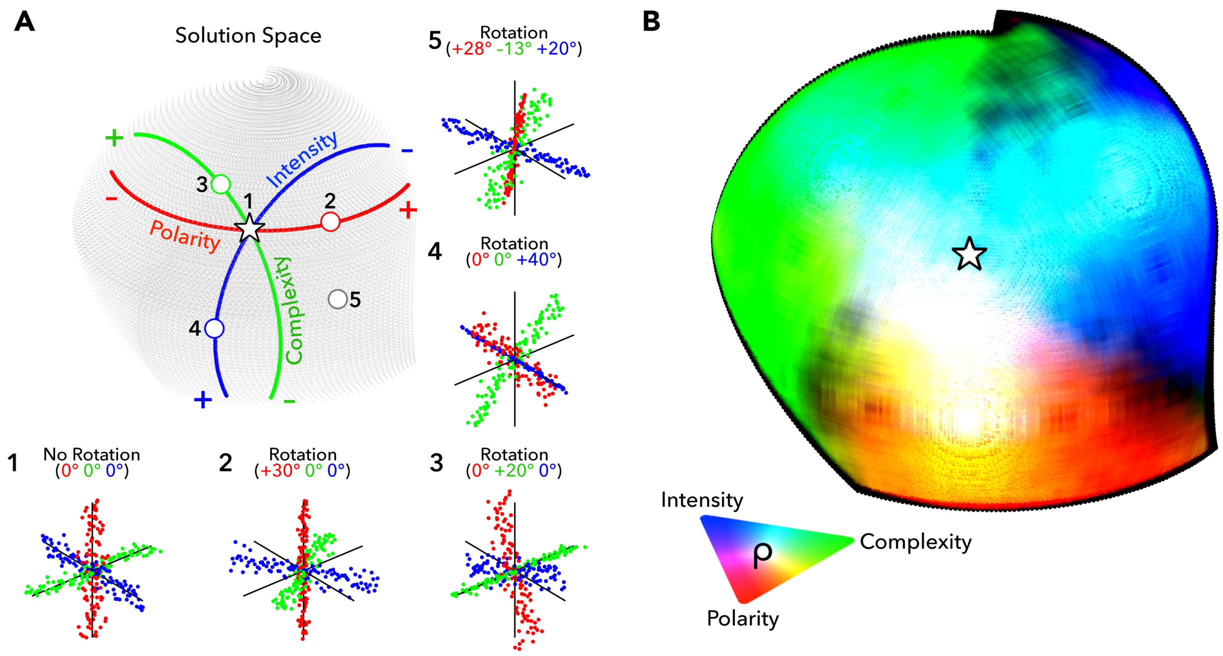


To test whether the rotated version of emotion dimensions explains the topographic organization of right TPJ, we systematically applied orthogonal rotations to polarity, complexity and intensity components, fitted each solution in brain activity and then estimated the magnitude of the obtained gradients. Panel **A** depicts the solution space: the pentagram (1) represents the solution determined by the unrotated principal components, whereas the red (2), green (3) and blue (4) round markers express the orthogonal rotation of two components while keeping fixed the other one. The grey round marker (5) maps a solution obtained by applying orthogonal rotations to the three axes. Scatter plots illustrate the transformations applied to the data for each of the points represented in the solution space. Panel **B** shows the effect size (i.e., Spearman's ρ) of the estimate of gradient for all the explored solutions. Magnitude of rotated polarity, complexity and intensity gradients is expressed by hue and brightness.

**Supplementary Figure 5. Brain areas representing the combination of emotion dimension gradients as revealed by searchlight analysis**

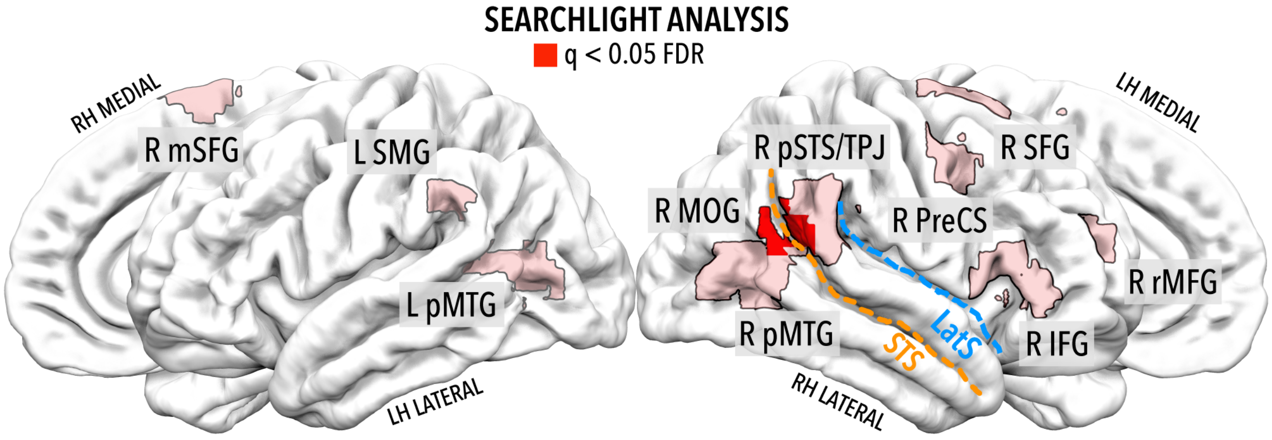


Shaded and outlined regions indicate voxels significantly encoding emotion ratings. In red results corrected using the False Discovery Rate procedure. Datasets for these results are available in the public repository. pSTS/TPJ = posterior part of the superior temporal sulcus/temporoparietal junction, pMTG = posterior middle temporal gyrus, preCS = precentral sulcus, IFG = inferior frontal gyrus, mSFG = medial superior frontal gyrus, SMG = supramarginal gyrus, rMFG = rostral middle frontal gyrus, MOG = middle occipital gyrus.

**Supplementary Figure 6. Brain areas representing individual emotion dimension gradients as revealed by searchlight analysis**

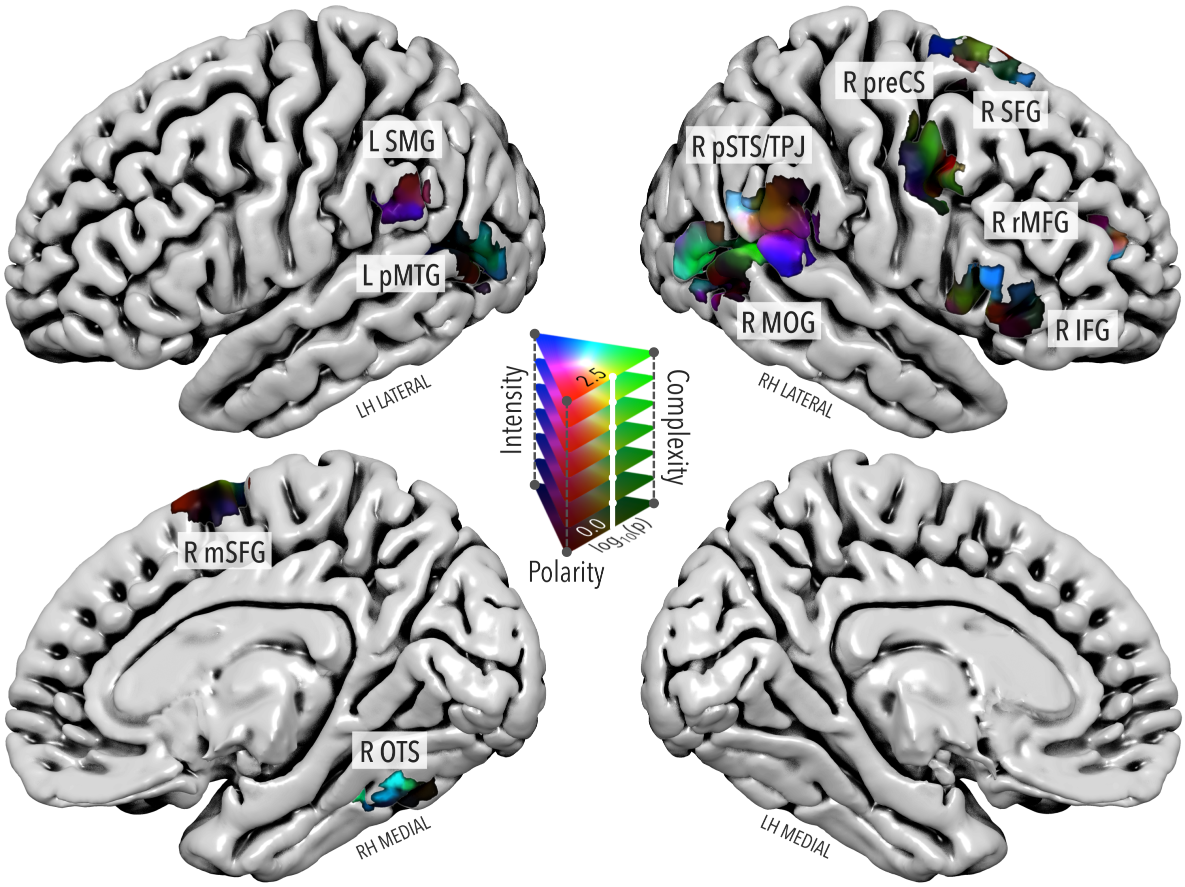


Results for the three separate searchlight analyses measuring the topographic arrangement of polarity (red channel), complexity (green channel) and intensity (blue channel). Color brightness relates to the log transformed p-value of the fitting of each component. Datasets for these results are available in the public repository. pSTS/TPJ = posterior part of the superior temporal sulcus/temporoparietal junction, pMTG = posterior middle temporal gyrus, preCS = precentral sulcus, IFG = inferior frontal gyrus, mSFG = medial superior frontal gyrus, SMG = supramarginal gyrus, rMFG = rostral middle frontal gyrus, MOG = middle occipital gyrus; OTS = occipitotemporal sulcus.

**Supplementary Figure 7. Emotion dimension gradients using unsmoothed fMRI data**

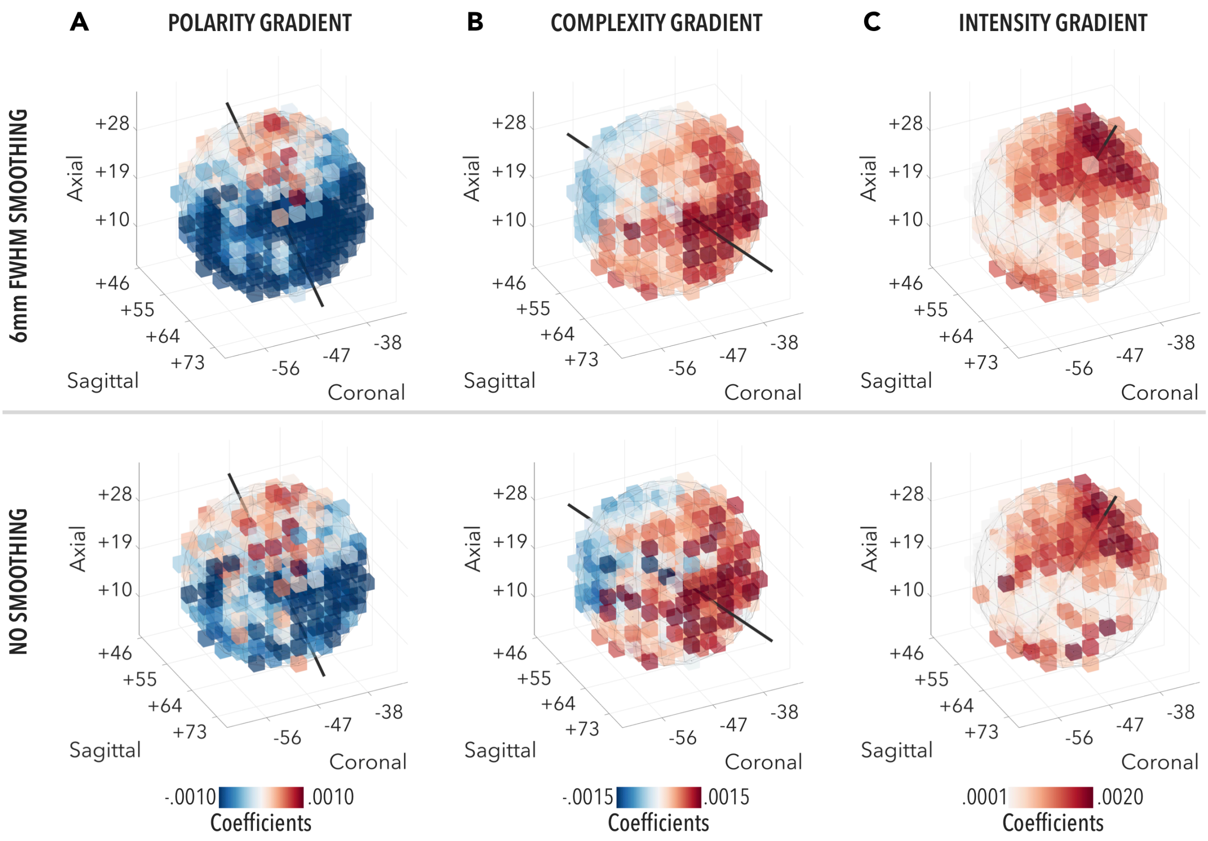


Uppermost row depicts β coefficients of emotion dimensions obtained when applying 6mm FWHM smoothing (3dBlurToFWHM). Lowermost row depicts β coefficients of emotion dimensions obtained without applying any spatial filtering. Panel **A**, **B** and **C** represent polarity, complexity and intensity gradients in right TPJ, respectively.

**Supplementary Figure 8. Subjective emotion rating model versus third person emotion attribution models**

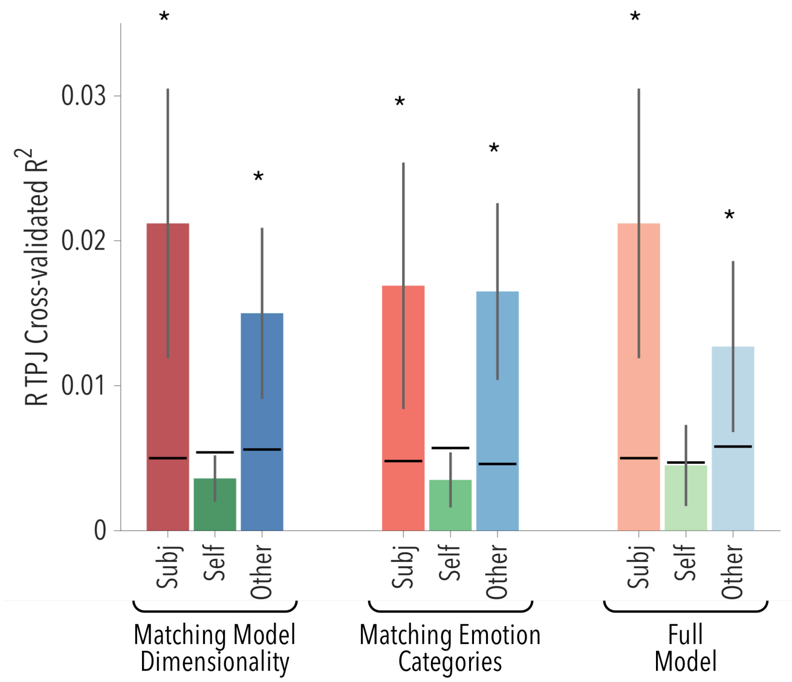


We measured the extent to which the two third-person emotion attribution models explained brain activity in right TPJ. We assessed the significance of fitting using three different procedures: (**A**) matching the dimensionality across models by selecting the first six principal components only; (**B**) matching the emotion categories in ratings, by performing PC analysis on the four basic emotions shared across models (i.e., happiness, fear, sadness and anger); (**C**) using the full model regardless of the dimensionality (i.e., six for our subjective emotion rating and 22 for the emotion attribution models). Results showed that only the subjective emotion rating model and the other-directed emotion attribution one significantly explained activity of right TPJ (**A**: subj $R^2 = 0.021$, $p < 0.002$; other $R^2 = 0.015$, $p < 0.002$; self $R^2 = 0.004$, $p = 0.269$. **B**: subj $R^2 = 0.017$, $p < 0.002$; other $R^2 = 0.016$, $p < 0.002$; self $R^2 = 0.003$, $p = 0.335$. **C**: subj $R^2 = 0.021$, $p < 0.002$; other $R^2 = 0.013$, $p < 0.002$; self $R^2 = 0.004$, $p = 0.078$). The subjective emotion rating and the other-directed emotion attribution model did not significantly differ in explaining activity of right TPJ ($p > 0.05$). The lower and upper noise ceiling bounds averaged across all the right TPJ voxels were $R^2 = 0.11$ and $R^2 = 0.20$. * denotes $p < 0.05$; error bar indicates standard error; bold horizontal line is the 95$^{th}$ percentile of the null distribution.

**Supplementary Figure 9. Preferred responses of distinct populations of voxels using non-negative matrix factorization**

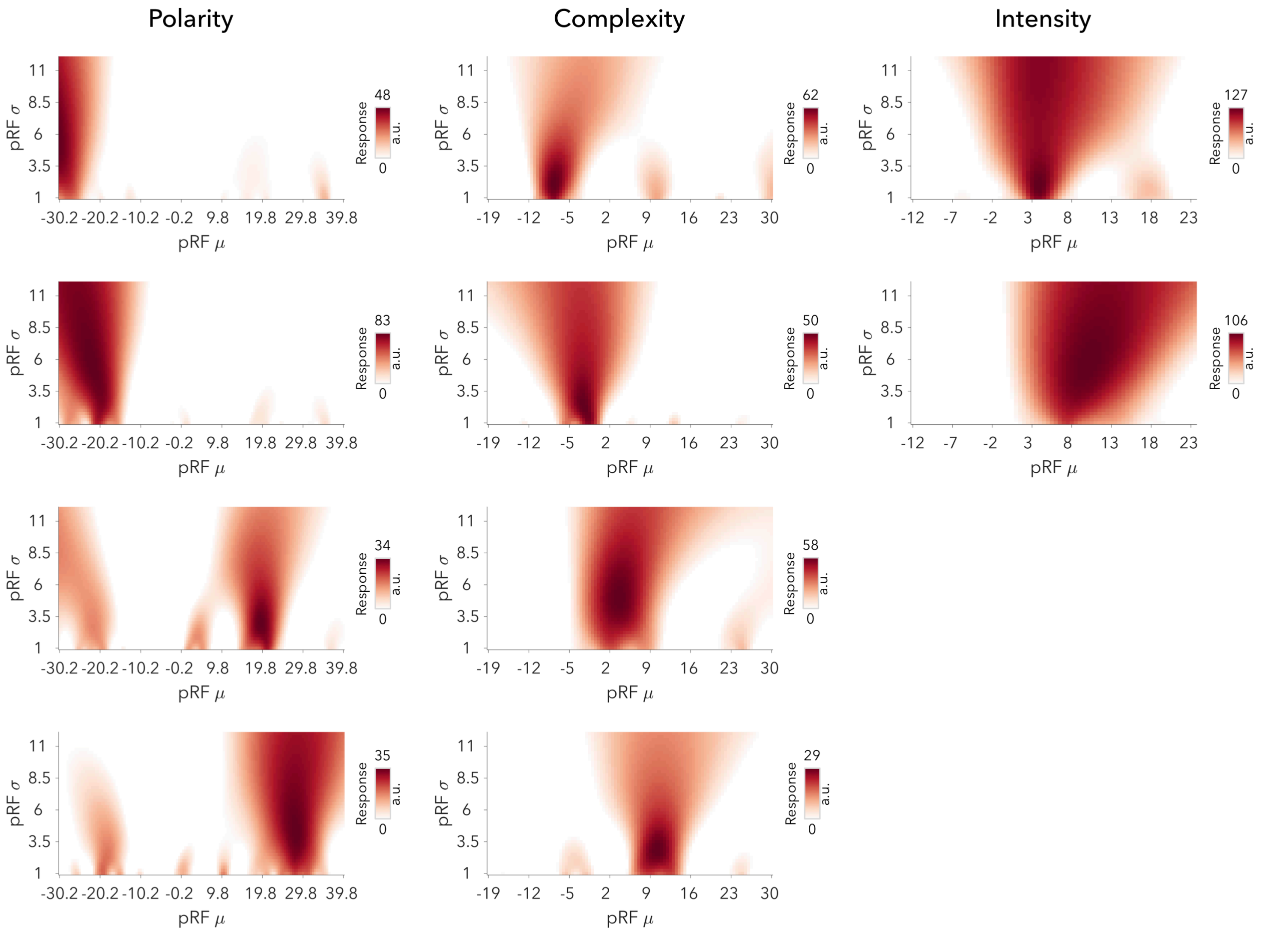


Prototypical responses of populations of voxels as function of affective states. We decomposed the pRF data (i.e., voxels t-values for all the explored μ and σ in the grid-search procedure) using non-negative matrix factorization. The figure depicts resulting components retaining at least 5% of the variance for polarity (i.e., first column), complexity (i.e., second column) and intensity (i.e., third column). Results highlight the existence of four distinct populations of voxels tuned to specific scores of polarity and complexity. Two populations represented distinct intensity values.

### Supplementary Figure 10. Fitting of low-level features

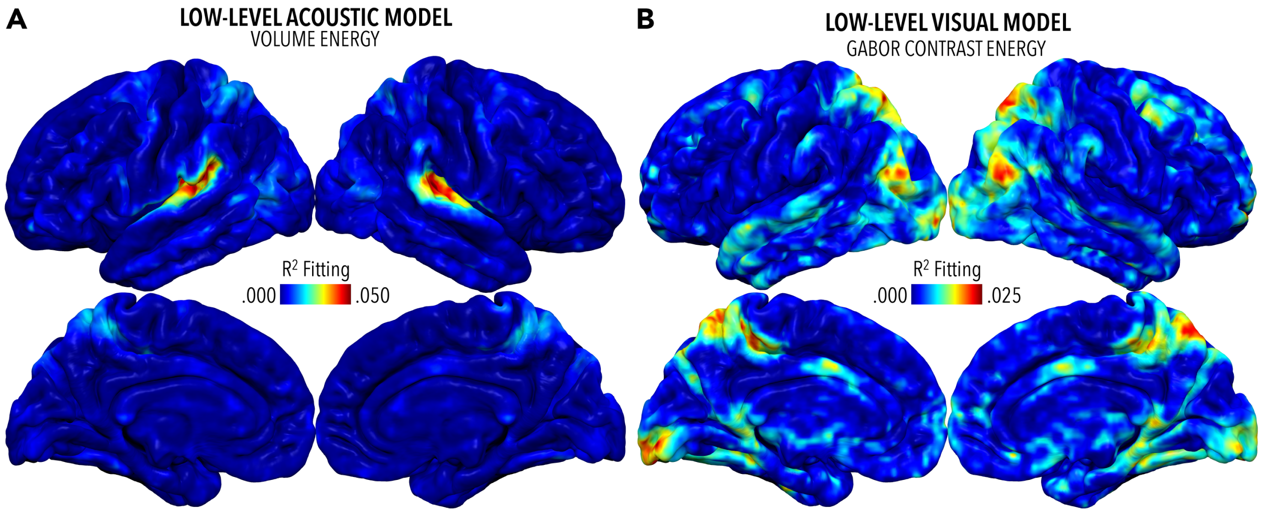


The fitting of low-level stimulus properties was estimated to verify the adequacy of adopted models. **A**. Peak of fitting for the volume energy model (i.e., RMS of the audio track) is located in primary auditory cortex. **B**. Gabor contrast energy (i.e., low and high spatial frequencies) of movie frames explained activity in primary visual cortex as well as in other associative areas (e.g., retrosplenial, parahippocampal and superior parietal cortex).

**Supplementary Figure 11. Reconstruction of the emotion dimensions from portrayed emotions**

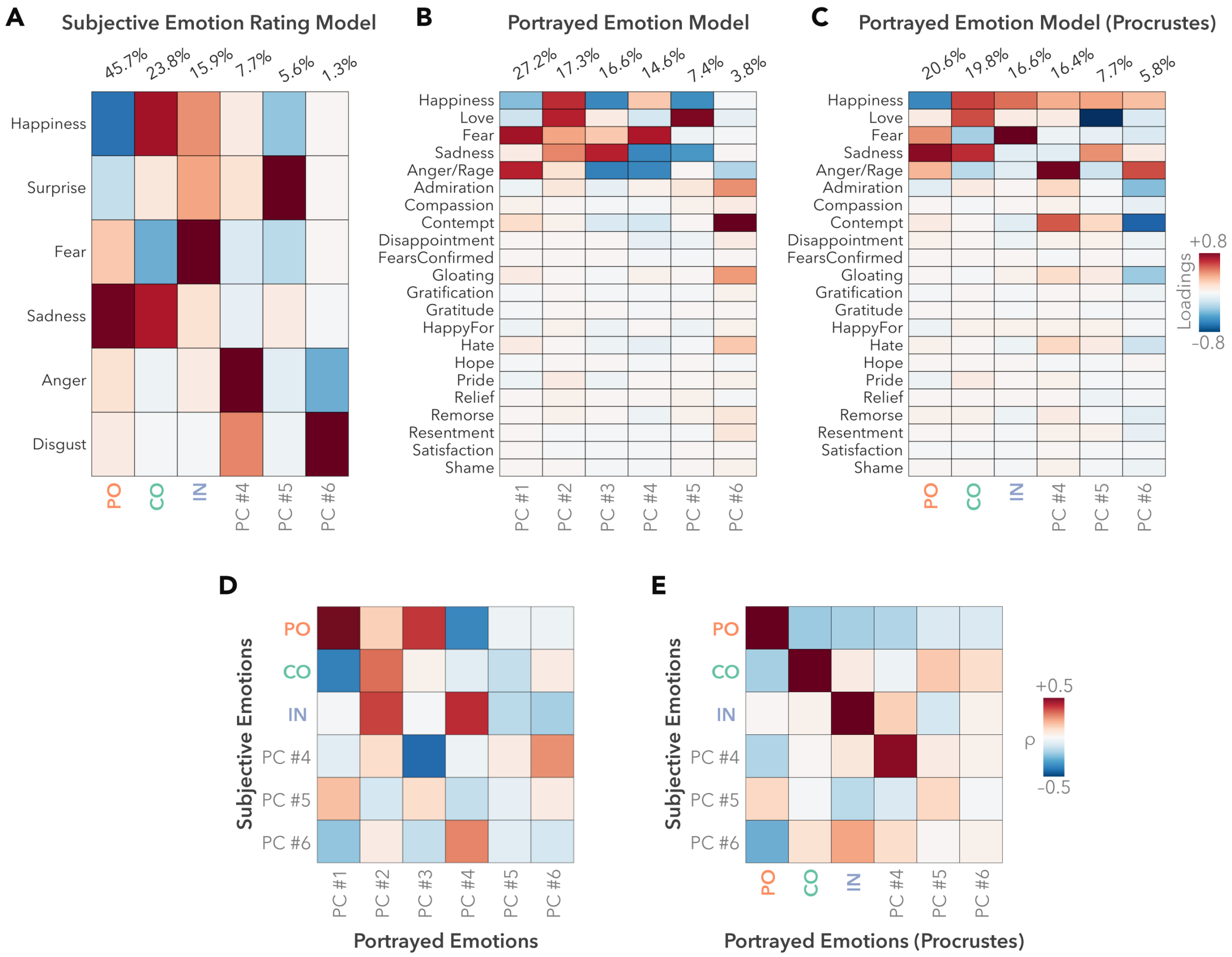


Panel **A** shows the six PCs obtained from subjective emotion ratings. Panel **B** depicts the output of PCA for Labs and colleagues[1] data. The first six dimensions represented ~85% of the explained variance. Panel **C** demonstrates that our polarity, complexity and intensity dimensions emerge from the portrayed emotion model after rotating PC scores using the procrustes criterion. Lowermost panels report the correlation between our original PCs and the unrotated (**D**) and rotated (**E**) version of components derived from portrayed emotions.

### Supplementary Figure 12. Real life versus Forrest Gump emotion transitions

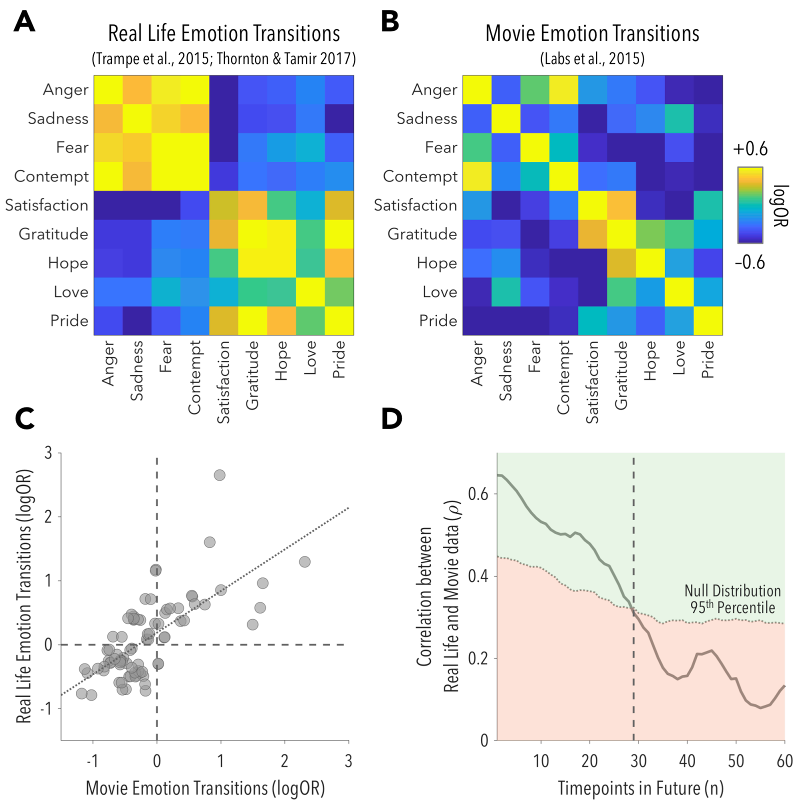


We analyzed data relative to study 3 of Thornton & Tamir 2017[6] and selected the emotion categories in common with Labs and coauthors 2015[1]. (**A**) Matrix depicting real life emotion transitions: each cell represents the log odds of a particular emotion transition. We built this matrix from an experience-sampling dataset of subjects reporting their affective state throughout the day[6]. (**B**) Matrix showing movie emotion transitions: each cell represents the log odds of a particular emotion transition during Forrest Gump. We built this matrix from the reports of portrayed emotions[1], similar to[6]. (**C**) Real life emotion transitions are significantly associated to the movie-based emotion transitions (Spearman's $\rho = 0.646$; $p = 0.001$). (**D**) We built a number of movie-based models, each measuring the likelihood of emotion transition between timepoint t and timepoint t+n in the future, with a maximum delay of 120 seconds (60 timepoints). These models were then correlated with (**A**) and results show that the real life model predicts emotion transitions in the movie up to 58 seconds.

### Supplementary Figure 13. fMRI data single-subject preprocessing

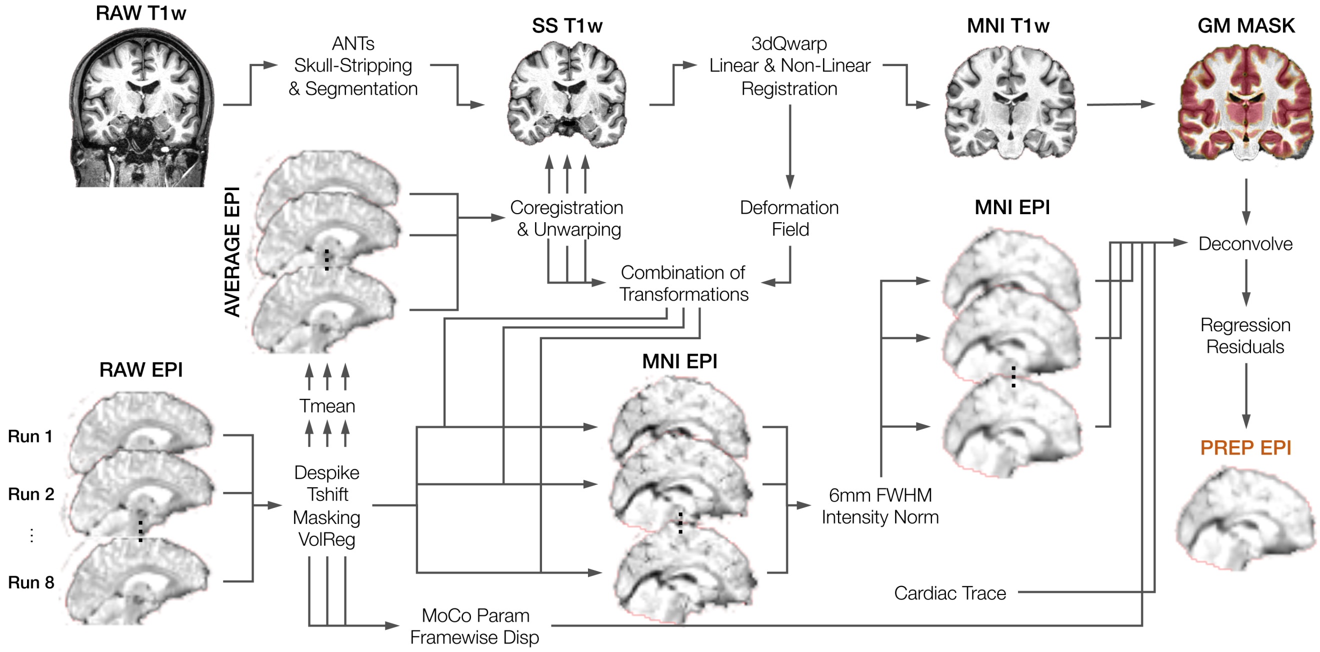


Preprocessing pipeline for structural and functional MRI data.

### Supplementary Figure 14. fMRI data group-level preprocessing

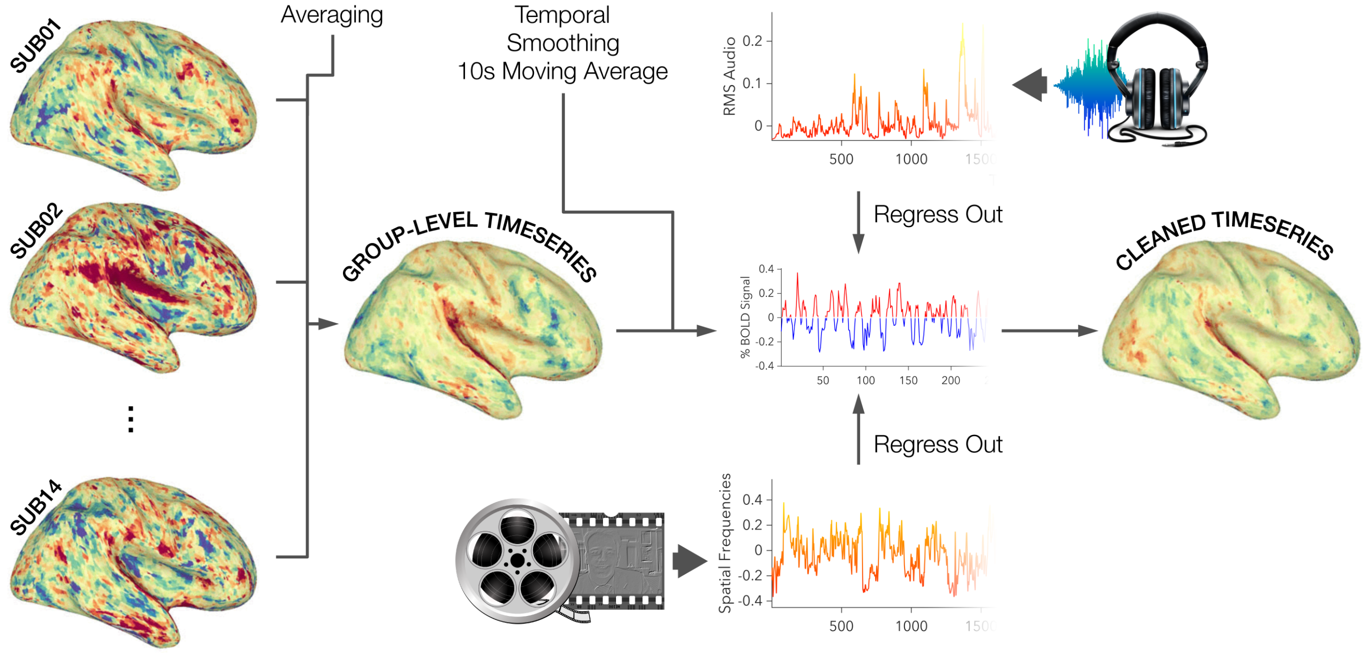


Single-subject preprocessed fMRI data were averaged to obtain group-level hemodynamic activity. For each voxel a windowing procedure was employed to temporally smooth data (moving average: 10s window). From the obtained aggregated and smoothed timeseries, the timecourse of low-level acoustic (i.e., volume energy - RMS of the signal) and visual (i.e., Gabor contrast energy for 0.5 and 8 cyc/deg spatial frequencies for each frame) movie features were regressed out, so as to mitigate the possible collinearity between emotion ratings and low-level psychophysical properties of the stimulus.

**Supplementary Figure 15. Comparison between emotion gradients and meta-analytic definition of right TPJ**

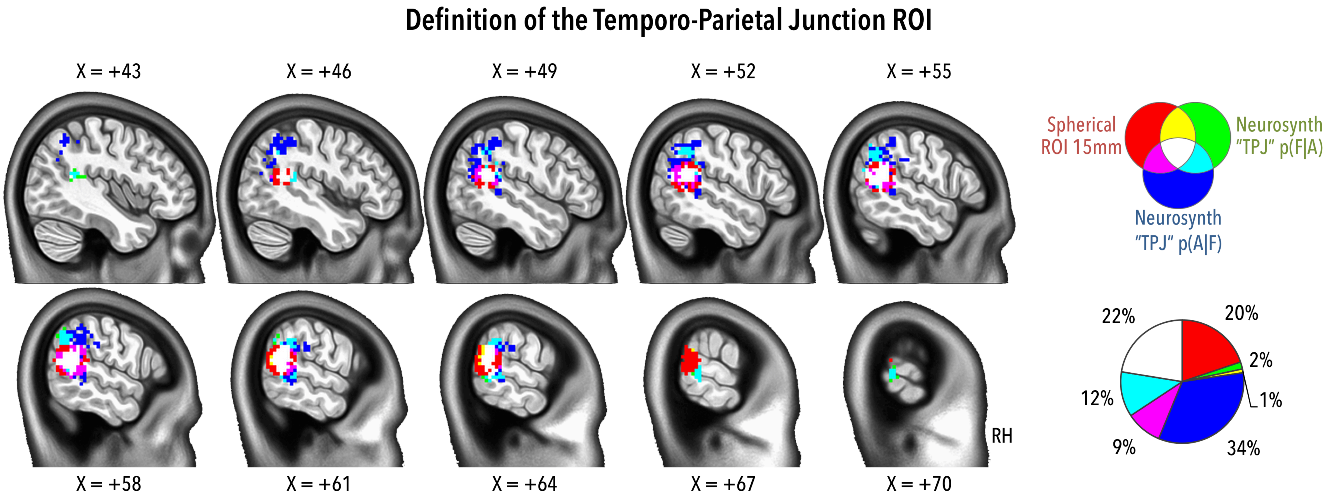


We obtained from the Neurosynth database (http://old.neurosynth.org/analyses/terms/tpj/) two meta-analytic maps representing a reliable estimate of the right TPJ size, against which we compared the volume of our spherical ROI. Neurosynth TPJ reverse inference map - p(F|A) - is represented in green, whereas the TPJ forward inference map - p(A|F) - is in blue and our spherical ROI (i.e., 15mm radius) is in red. Pie chart represents the percentage of volume related to our ROI, the two meta-analytic maps and their overlap.

**Supplementary Figure 16. Regions associated to the direction of portrayed emotions**

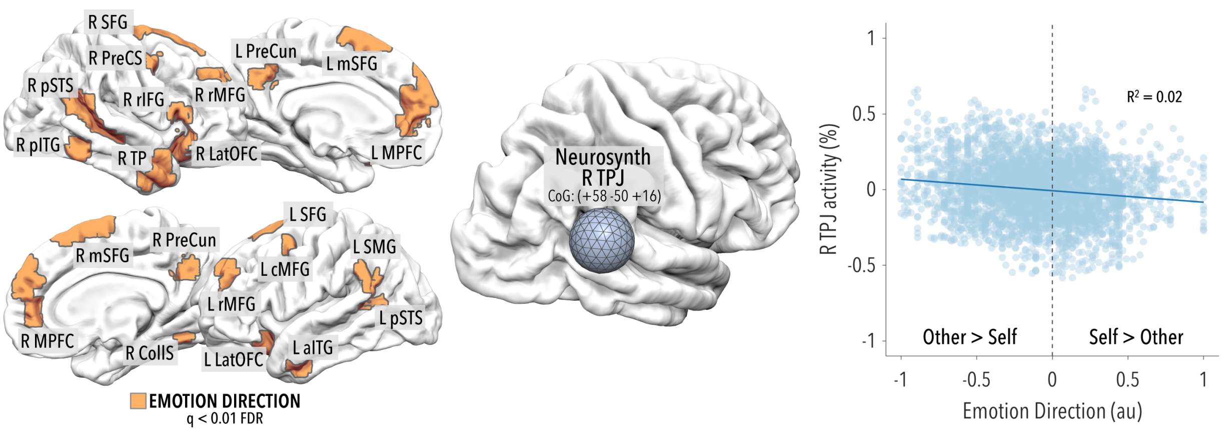


We performed a voxel-wise encoding of the direction of portrayed emotions[1] on group-averaged BOLD signal. The higher the BOLD of right TPJ, the more raters labeled emotions as other-directed (right TPJ peak $R^2$: 0.04; right TPJ average $R^2$: 0.02). Significant associations ($p < 0.01$ FDR corrected) between emotion direction and BOLD signal were also found in other brain regions of the Theory of Mind, empathy and emotion processing networks, closely resembling the pattern found by Hanke and colleagues[27]. Datasets for these results are available in the public repository. CoG = center of gravity; pSTS = posterior part of the superior temporal sulcus, preCS = precentral sulcus, IFG = inferior frontal gyrus, mSFG = medial superior frontal gyrus, SMG = supramarginal gyrus, rMFG = rostral middle frontal gyrus, TP = temporal pole, CollS = collateral sulcus, aITG = anterior inferior temporal gyrus, cMFG = caudal middle frontal gyrus, SFG = superior frontal gyrus, LatOFC = lateral orbitofrontal cortex, MPFC = middle prefrontal cortex, PreCun = precuneus, pITG = posterior inferior temporal gyrus.

**Supplementary Figure 17. Right TPJ emotion dimension gradients do not depend on low-level acoustic and visual features**

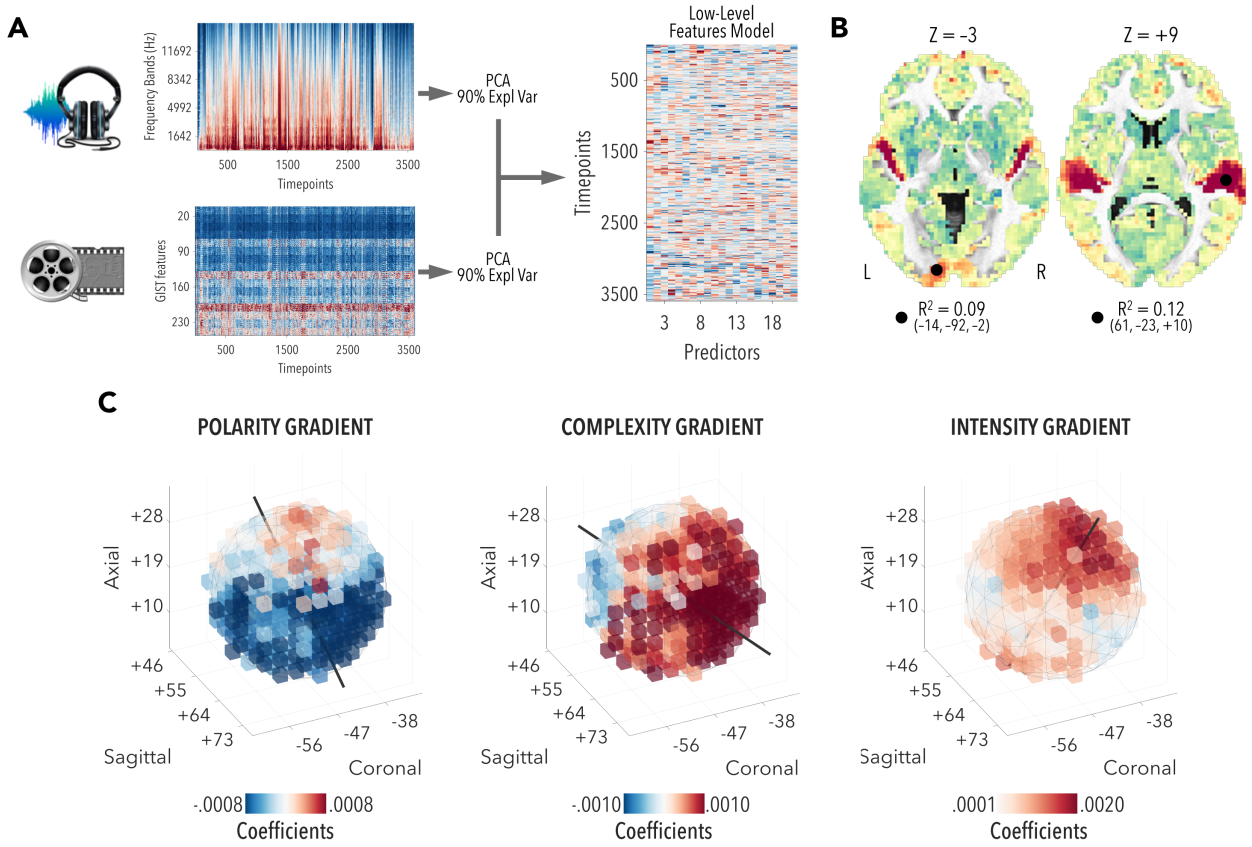


**Panel A** represents the 21 PCs used to model acoustic and visual low-level properties of the Forrest Gump movie derived from power spectral features and GIST descriptors. Using this model, we have more than doubled the explained variance in sensory cortical areas (12% in Heschl's gyrus and 9% in pericalcarine cortex; **B**), as compared to RMS and contrast energy models (Supplementary Figure 10). Of note, upper and lower noise ceiling bounds for the highest $R^2$ voxels were 0.268-0.172 in primary auditory cortex and 0.412-0.330 in early visual cortex. These numbers suggest that our 21 PCs model explains up to 70% and 27% of brain activity within these regions. Most importantly, we found that right TPJ emotion dimensions gradient were not affected by regressing out low-level properties from BOLD signal (**C**): polarity ($\rho$ = 0.258, p-value = 0.031, 95% CI: 0.252 to 0.264), complexity ($\rho$ = 0.261, p-value = 0.013, 95% CI: 0.254 to 0.267) and intensity ($\rho$ = 0.270, p-value = 0.016, 95% CI: 0.264 to 0.277).

### Supplementary Tables

#### Supplementary Table 1. Brain regions encoding emotion ratings

| | Cluster Size | Peak | | | CoG | | |
|---|---|---|---|---|---|---|---|
| | | x | y | z | x | y | z |
| R pSTS/TPJ | 343 | 61.5 | -40.5 | 19.5 | 55.9 | -54.1 | 16 |
| L pMTG | 96 | -49.5 | -52.5 | 10.5 | -51 | -63.8 | 8.4 |
| R preCS | 95 | 49.5 | 1.5 | 55.5 | 48.7 | 3.1 | 44.6 |
| R IFG | 42 | 55.5 | 22.5 | 10.5 | 50 | 22.1 | 5.6 |
| R mSFG | 30 | 1.5 | -4.5 | 70.5 | 6 | -1.3 | 68.8 |
| R OTS | 28 | 49.5 | -46.5 | -19.5 | 46.7 | -44.6 | -20.7 |
| L SMG | 20 | -55.5 | -40.5 | 31.5 | -60.8 | -41 | 32.6 |
| R IFG | 15 | 55.5 | 31.5 | -1.5 | 53.2 | 33.1 | 1.3 |
| R mSFG | 15 | 13.5 | 16.5 | 61.5 | 14 | 14.4 | 61.1 |
| R mSFG | 15 | 16.5 | 1.5 | 67.5 | 14.8 | 3 | 69 |
| R rMFG | 12 | 22.5 | 52.5 | 19.5 | 23.7 | 53 | 17.5 |

Table showing regions significantly associated to emotion ratings ($q < 0.01$; minimum cluster size > 10 voxels). Voxel size = 3 mm isotropic; CoG = center of gravity; pSTS/TPJ = posterior part of the superior temporal sulcus/temporoparietal junction, pMTG = posterior middle temporal gyrus, preCS = precentral sulcus, IFG = inferior frontal gyrus, mSFG = medial superior frontal gyrus, OTS = occipitotemporal sulcus, SMG = supramarginal gyrus, rMFG = rostral middle frontal gyrus.

### Supplementary Table 2. Emotion gradients in TPJ

| | Radius (mm) | Emotion Dimensions | | Basic Emotions | |
|---|---|---|---|---|---|
| | | ρ | p-value | ρ | p-value |
| Right TPJ | 9 | **0.316** | **0.037** | 0.286 | 0.076 |
| | 12 | **0.375** | **0.001** | **0.326** | **0.024** |
| | 15 | **0.399** | **<0.001** | **0.352** | **0.004** |
| | 18 | **0.387** | **<0.001** | **0.332** | **0.006** |
| | 21 | **0.372** | **<0.001** | **0.319** | **0.007** |
| | 24 | **0.342** | **0.001** | **0.299** | **0.012** |
| | 27 | **0.292** | **0.003** | **0.266** | **0.021** |
| Left TPJ | 15 | 0.251 | 0.144 | 0.208 | 0.356 |

To identify the patch of cortex with the highest significant association between anatomical and functional distance, we started from the reverse inference peak for the term "TPJ" in the NeuroSynth database. We then created a set of spherical ROIs having as center of gravity this peak and with radius ranging from 9 to 27 mm. For each ROI, we tested the relationship between anatomical and functional distance using the procedure detailed in the main manuscript and depicted in Supplementary Figure 2. The procedure was performed using either the three emotion dimensions or the four basic emotions stable across all subjects. Results demonstrated that within a 15 mm radius ROI, relative spatial arrangement and functional features of right TPJ were significantly and maximally correlated either considering the basic emotion model or the emotion dimensions one. Moreover, we included a 15 mm ROI centered at the left TPJ as control region (Neurosynth definition). TPJ = Temporoparietal junction.

**Supplementary Table 3. Emotion gradients in TPJ relative to emotion dimensions or basic emotions**

| | Radius | Polarity | | Complexity | | Intensity | | PC #4 | | PC #5 | | PC #6 | |
|---|---|---|---|---|---|---|---|---|---|---|---|---|---|
| | (mm) | ρ | p-value | ρ | p-value | ρ | p-value | ρ | p-value | ρ | p-value | ρ | p-value |
| Right TPJ | 15 | **0.241** | **0.041** | **0.271** | **0.013** | **0.229** | **0.049** | 0.044 | 0.975 | 0.239 | 0.052 | 0.114 | 0.598 |
| Left TPJ | 15 | 0.132 | 0.354 | 0.157 | 0.222 | 0.149 | 0.257 | 0.088 | 0.643 | 0.049 | 0.889 | 0.171 | 0.169 |

| | Radius | Happiness | | Surprise | | Fear | | Sadness | | Anger | | Disgust | |
|---|---|---|---|---|---|---|---|---|---|---|---|---|---|
| | (mm) | ρ | p-value | ρ | p-value | ρ | p-value | ρ | p-value | ρ | p-value | ρ | p-value |
| Right TPJ | 15 | **0.275** | **0.013** | 0.202 | 0.112 | 0.197 | 0.091 | 0.182 | 0.160 | 0.141 | 0.379 | 0.097 | 0.724 |
| Left TPJ | 15 | 0.158 | 0.216 | 0.028 | 0.964 | 0.142 | 0.293 | 0.156 | 0.213 | 0.073 | 0.733 | 0.163 | 0.179 |

For each individual emotion dimension and basic emotion, we tested the existence of a gradient-like organization in a spherical ROI (15 mm radius) located within the TPJ region (Neurosynth definition). Results for emotion dimensions and basic emotions consistent across all subjects are reported in black (see the Agreement across subjects of the six basic emotions and Agreement across subjects of the emotion dimensions sections). Significant results are marked with bold. TPJ = Temporoparietal junction.

**Supplementary Table 4. Single-subject emotion dimension gradients in right TPJ**

| Sub ID | Gender | Polarity | | Complexity | | Intensity | |
|---|---|---|---|---|---|---|---|
| | | ρ | p-value | ρ | p-value | ρ | p-value |
| 02 | M | **0.464** | **0.019** | **0.446** | **0.002** | 0.411 | 0.071 |
| 03 | F | **0.349** | **0.013** | **0.362** | **0.029** | **0.521** | **<0.001** |
| 04 | F | **0.607** | **0.002** | **0.465** | **<0.001** | **0.423** | **0.007** |
| 05 | M | 0.074 | 0.244 | 0.199 | 0.229 | -0.004 | 0.508 |
| 06 | M | **0.660** | **0.003** | **0.639** | **0.001** | **0.480** | **<0.001** |
| 09 | M | **0.354** | **0.009** | **0.313** | **0.017** | 0.245 | 0.056 |
| 10 | F | **0.423** | **0.009** | **0.347** | **0.038** | **0.263** | **0.045** |
| 14 | F | 0.306 | 0.098 | **0.495** | **<0.001** | 0.231 | 0.117 |
| 15 | M | **0.232** | **0.044** | 0.183 | 0.098 | 0.151 | 0.182 |
| 16 | M | **0.380** | **0.006** | 0.256 | 0.103 | **0.561** | **<0.001** |
| 17 | M | **0.291** | **0.015** | 0.188 | 0.081 | **0.407** | **0.041** |
| 18 | M | 0.154 | 0.210 | **0.448** | **0.008** | **0.369** | **0.017** |
| 19 | F | **0.619** | **0.001** | **0.485** | **0.015** | **0.559** | **<0.001** |
| 20 | F | **0.594** | **<0.001** | **0.425** | **0.001** | **0.580** | **<0.001** |

We tested the consistency of emotion dimension gradients in right TPJ using single-subject data. Firstly, preprocessed fMRI single-subject timeseries were smoothed in time (10s moving average window) and cleaned from low-level visual and acoustic features of the movie, as in the group-level analysis pipeline. Subsequently, we performed an encoding analysis using the behavioral ratings and obtained β values for polarity, complexity and intensity. Afterwards, we measured the relationship between single-subject maps and those obtained from group-level analysis using Spearman's ρ coefficient. To measure the statistical significance of these associations, we employed a surrogate-based approach by generating 1,000 emotion dimension encoding models using the IAAFT procedure, as described in the Methods section. Bold values represent significant associations between single-subject and group-level gradients.

**Supplementary Table 5. Topographic organization of portrayed emotions in right TPJ**

| PC | Other-directed model | | Other-directed CCA | |
|---|---|---|---|---|
| # | ρ | p-value | ρ | p-value |
| 1 | 0.136 | 0.404 | **0.221** | **0.036** |
| 2 | 0.111 | 0.594 | 0.150 | 0.384 |
| 3 | 0.119 | 0.554 | 0.207 | 0.092 |
| 4 | 0.081 | 0.844 | 0.062 | 0.897 |
| 5 | 0.157 | 0.276 | 0.101 | 0.684 |
| 6 | 0.118 | 0.533 | 0.107 | 0.664 |
| 7 | 0.147 | 0.363 | | |
| 8 | 0.105 | 0.676 | | |
| 9 | 0.178 | 0.234 | | |
| 10 | 0.190 | 0.154 | | |
| 11 | 0.124 | 0.546 | | |
| 12 | 0.177 | 0.200 | | |
| 13 | 0.158 | 0.270 | | |
| 14 | 0.124 | 0.511 | | |
| 15 | 0.154 | 0.340 | | |
| 16 | 0.115 | 0.587 | | |
| 17 | 0.094 | 0.735 | | |
| 18 | **0.290** | **0.004** | | |
| 19 | 0.101 | 0.688 | | |
| 20 | 0.105 | 0.669 | | |
| 21 | 0.130 | 0.485 | | |
| 22 | 0.124 | 0.519 | | |

We tested right TPJ topography for the other-directed emotion attribution PCs. None of the first six components retained a topographical organization in this region. Only the $18^{th}$ PC, explaining the 0.3% of the variance, appeared to be encoded in a gradient-like manner. However, the pattern associated to this component was also collinear with activity evoked by polarity ($\rho = 0.494$) and intensity ($\rho = 0.475$) dimensions. Moreover, using CCA (canonical correlation analysis) we transformed the 22-dimensional space defined by the other-directed model to match our subjective reports. Noteworthy, when fitting the aligned components into right TPJ activity, only the first PC (i.e., reconstructed polarity) was represented through a gradient. PC = principal component.

**Supplementary Table 6. Topographies in right TPJ considering spatial smoothing and cortical folding**

| | Radius (mm) | Polarity | | Complexity | | Intensity | | PC #4 | | PC #5 | | PC #6 | |
|---|---|---|---|---|---|---|---|---|---|---|---|---|---|
| | | ρ | p-value | ρ | p-value | ρ | p-value | ρ | p-value | ρ | p-value | ρ | p-value |
| Right TPJ | 15 | **0.167** | **0.033** | **0.186** | **0.010** | **0.184** | **0.010** | 0.014 | 0.996 | **0.194** | **0.018** | 0.076 | 0.584 |

| | Radius (mm) | Polarity | | Complexity | | Intensity | | PC #4 | | PC #5 | | PC #6 | |
|---|---|---|---|---|---|---|---|---|---|---|---|---|---|
| | | ρ | p-value | ρ | p-value | ρ | p-value | ρ | p-value | ρ | p-value | ρ | p-value |
| Right TPJ | 15 | **0.248** | **0.026** | **0.314** | **0.001** | **0.249** | **0.013** | 0.012 | 0.961 | 0.130 | 0.323 | 0.083 | 0.577 |

Table showing the robustness of emotion dimension gradients in right TPJ using unsmoothed data. The first row regards the evaluation of gradients in the unfiltered volumetric space. The second row refers to the results of the same analysis conducted with unfiltered data into surface space, using the Dijkstra algorithm. PC = principal component; TPJ = Temporoparietal junction.
